# O-GlcNAcylation promotes YTHDF1 cytosolic localization and colorectal cancer tumorigenesis

**DOI:** 10.1101/2022.11.21.517456

**Authors:** Jie Li, Muhammad Ahmad, Lei Sang, Yahui Zhan, Yibo Wang, Yonghong Yan, Yue Liu, Weixiao Mi, Mei Lu, Yu Dai, Rou Zhang, Meng-Qiu Dong, Yun-Gui Yang, Xiaohui Wang, Jianwei Sun, Jing Li

## Abstract

O-linked N-acetylglucosamine (O-GlcNAc) is an emerging post-translation modification that couples metabolism with cellular signal transduction by crosstalking with phosphorylation and ubiquitination to orchestrate various biological processes. Herein we show that it modifies the N^6^-methyladenosine (m^6^A)-mRNA reader YTHDF1 and fine-tunes its nuclear translocation by the exportin protein Crm1. First we present evidence that YTHDF1 interacts with the sole O-GlcNAc transferase (OGT). Second, we verified the YTHDF1 O-GlcNAcylation sites to be Ser196/Ser197/Ser198, as described in previous numerous chemoproteomic studies. Then we constructed the O-GlcNAc-deficient YTHDF1-S196AS197FS198A (AFA) mutants, which significantly attentuated O-GlcNAc signals. Moreover, we revealed that YTHDF1 is a nucleocytoplasmic protein, whose nuclear export is mediated by Crm1. Furthermore, O-GlcNAcylation increases the cytosolic portion of YTHDF1 by enhancing binding with Crm1, thus upregulating the downstream target (e.g. c-Myc) expression. Molecular dynamics simulations suggest that O-GlcNAcylation at S197 might promote the binding between the nuclear export signal motif and Crm1 through increasing hydrogen bonding. Mouse xenograft assays further demonstrate that YTHDF1-AFA mutants decreased the colon cancer mass and size via decreasing c-Myc expression. In sum, we found that YTHDF1 is a nucleocytoplasmic protein, whose cytosolic localization is dependent on O-GlcNAc modification. We propose that the OGT-YTHDF1-c-Myc axis might underlie colorectal cancer tumorigenesis.

## Introduction

The N^6^-methyladenosine (m^6^A) modification is quite abundant on internal mRNAs and its function and regulation has caught a wave of intense investigations (1,2). Its numerous writers, erasers and readers are under stringent control (3), and one of the readers is YTH domain family 1 (YTHDF1) (4). YTHDF1 promotes translation efficiency during arsenite recovery (5). YTHDF1 enhances translation in adult mouse dorsal root ganglions during injury recovery and augments axonal regeneration (6). YTHDF1 fuels translation upon neuronal stimuli, which is conducive to learning and memory (7). YTHDF1 also recognizes m^6^A-marked lysosomal protease mRNAs, thus mediating the decay of neoantigens and bolstering tumor suppressive immunotherapy(8). Recently, YTHDF1 and YTHDF3 are also found to promote stress granule formation, as m^6^A mRNAs are found to be enriched in stress granules (9).

The interconnection between m^6^A mRNA and cancer are being revealed (10-12), as m^6^A takes part in many aspects of tumor biology: cancer stem cell, tumor cell proliferation or oncogene expression. YTHDF1, in particular, has been found to be at the nexus of multiple tumorigenic pathways. YTHDF1 binds the m^6^A modified mRNA of c-Myc, whose enhanced translation would promote glycolysis and cancer cell proliferation (13). In non-small cell lung cancer, YTHDF1 upregulates the translation efficiency of CDK2, CDK cyclin D1, and YTHDF1 is also elevated in high-altitude people, possibly through the hypoxia Keap1-Nrf2-AKR1C1 pathway (14). In gastric cancer, YTHDF1 enhances the expression of frizzled 7 (FZD7), a key Wnt receptor that would hyper-activate the Wnt/β-catenin pathway (15). In ovarian cancer, YTHDF1 promotes the translation of Eukaryotic Translation Initiation Factor 3 Subunit C (EIF3C), a component of the protein translation initiation factor EIF3 complex (16). In cervical cancer, YTHDF1 elevates the translation of hexokinase 2 (HK2) via binding with its 3’-UTR, thus promoting the Warburg effect (17). All these results suggest that YTHDF1 binds with its targets via m^6^A mRNA, and plays a fundamental role during human carcinogenesis.

Investigations show that some of the m^6^A regulators are subject to post-translational modifications (PTMs). YTHDF2, another m^6^A reader that mediates mRNA decay (18), is subject to SUMOylation at K571 upon hypoxia stress (19). SUMOylation would alter the binding affinity of YTHDF2 with m^6^A, thus deregulating the downstream target genes, leading to lung cancer progression (19). An m^6^A writer, Methyltransferase-like 3 (METTL3), is modified by lactylation at its zinc-finger domain, which changes its RNA capturing capacity, and regulates immunosuppression of tumor-infiltrating myeloid cells (20). Mettl3 is also acetylated, which regulates its localization and cancer metastasis (21).

The O-linked N-acetylglucosamine (O-GlcNAc) glycosylation is one PTM that occurs intracellularly (22) (23). Functioning as a rheostat to environmental stress or cellular nutrient status, O-GlcNAc monitors transcription, neural development, cell cycle and stress response (22) (23). However, whether it plays a role in m^6^A regulation has remained enigmatic. Historically O-GlcNAc studies have been strenuous due to technical impediment. Recent years have witnessed the combined methodology of chemoenzymatic labeling, bioorthogonal conjugation and ETD mass spectrometry, which have smoothened the way for biological investigations. Previously, an isotope-tagged cleavable linker together with chemoenzymatic labeling screen has identified the O-GlcNAc sites of YTHDF1 to be S196 and S198 (24). A second enrichment strategy using Gal labeling followed by chemical oxidation points the YTHDF1 O-GlcNAcylation region to be Ser196-198 (25). In another isotope targeted glycoproteomic study in T cells, YTHDF1 O-GlcNAcyation occurs on several residues, including Ser196, Ser197 and Ser198 (26). In this manuscript, we first confirmed that YTHDF1 O-GlcNAcylation occurs on Ser196Ser197Ser198. Then we found that YTHDF1 is a nucleocytoplasmic protein with exportin 1 (CRM1) mediating its cytoplasmic shuttling. We further presented evidence that O-GlcNAcylation promotes YTHDF1 cytosolic localization, thus enhancing downstream target expression, such as c-Myc. Our results were further correlated with TCGA analysis combined with mouse xenograft models. Our data highlight the signification of glycosylation in m^6^A regulation and tumorigenesis.

## Results

### YTHDF1 is O-GlcNAcylated at Ser196 Ser197 Ser198

As YTHDF1 has reproducibly been identified in O-GlcNAc profiling screens (27-29), we first assessed the binding affinity between YTHDF1 and the sole O-GlcNAc writer-OGT. 293T cells were transfected with Flag-YTHDF1 and HA-OGT plasmids, and the two overproduced proteins coimmunoprecipitate (coIP) (Fig. 1A). When the endogenous proteins were examined, YTHDF1 proteins were also present in the anti-OGT immunoprecipitates (Fig. 1B). Then pulldown assays were utilized to evaluate the physical association. 293T cells were transfected with HA-OGT, and the cell lysates were incubated with recombinant GST-YTHDF1 proteins. GST-YTHDF1 pulled-down overproduced OGT proteins (Fig. 1C). When pulldown assays were carried out between recombinant OGT and YTHDF1, again GST-YTHDF1 pulled-down His-OGT (Fig. 1D), suggesting that OGT and YTHDF1 directly interact *in vivo* and *in vitro*.

**Fig. 1.**
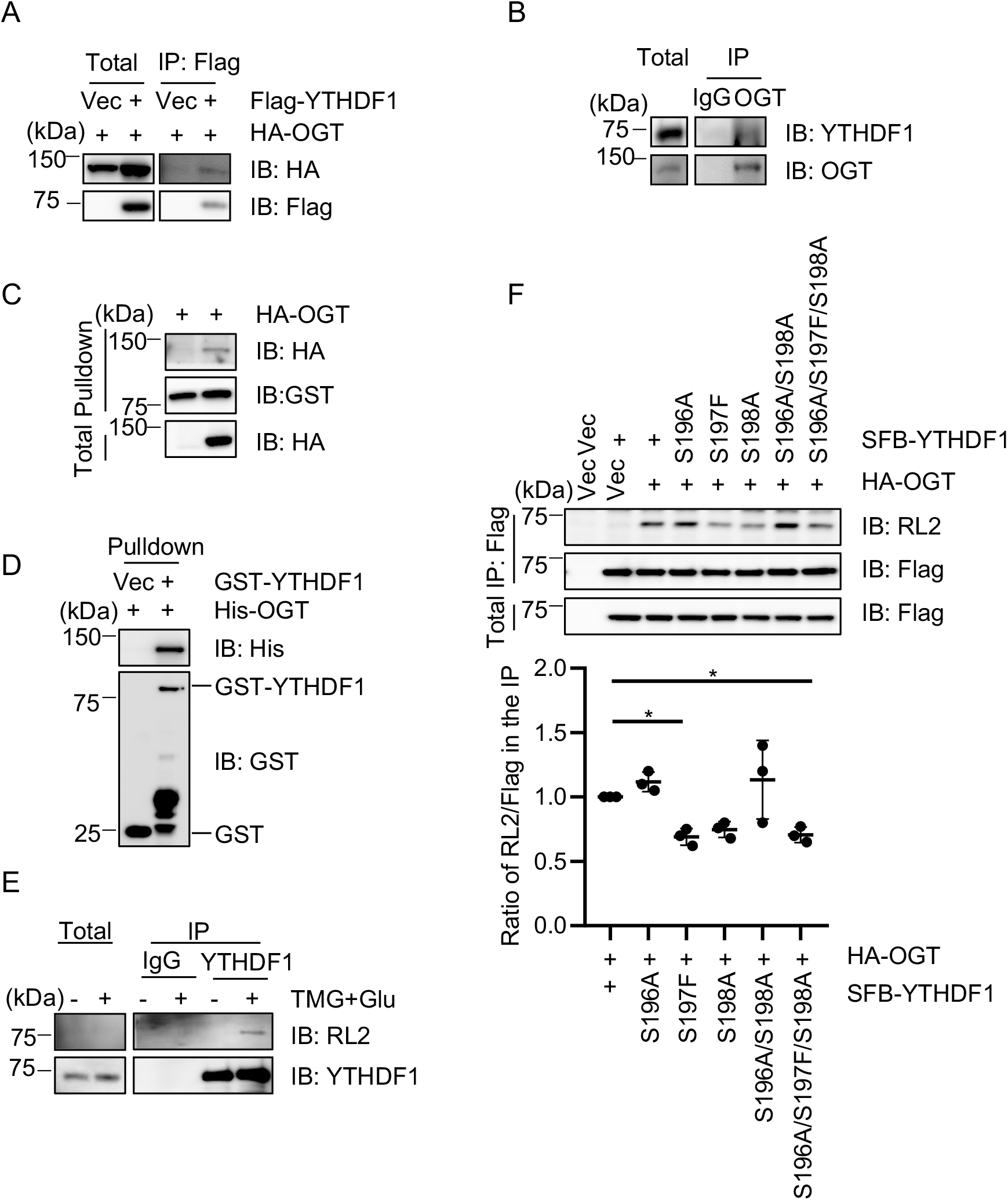
YTHDF1 is O-GlcNAcylated at Ser196 Ser197 Ser198. (A)293T cells were transfected with Flag-YTHDF1 and HA-OGT. The cell lysates were subject to immunoprecipitation and immunoblotting with the antibodies indicated. (B) HeLa cell lysates were immunoprecipitated with anti-OGT antibodies and immunoblotted with the indicated antibodies. (C) 293T cells were transfected with HA-OGT and the cell lysates were incubated with recombinant GST-YTHDF1. (D) Recombinant His-OGT and GST-YTHDF1 proteins were incubated and subject to pulldown assays. (E) Cells were treated with the OGA inhibitor Thiamet-G (TMG) and glucose to enrich for O-GlcNAcylation as described previously (30). Then the cell lysates were immunoprecipitated with anti-YTHDF1 antibodies and immunoblotted with anti-O-GlcNAc RL2 antibodies. (F) YTHDF1-S196A, S197F, S198A, -S196 (AFA) mutants were constructed and the cells were transfected with HA-OGT together with SFB-YTHDF1-WT, and -AFA mutants. The anti-Flag immunoprecipitates were immunoblotted with RL2 antibodies.

Then we assessed the O-GlcNAcylation of YTHDF1. 293T cells were enriched for O-GlcNAc by supplementing the media with glucose and Thiamet-G (TMG, the OGA inhibitor) (TMG + Glu) as previously described (30,31). The endogenous YTHDF1 proteins were IPed from the lysates, and RL2 antibodies detected a crisp band upon O-GlcNAc enrichment (Fig. 1E), suggesting that YTHDF1 is indeed O-GlcNAcylated. We decided to mutate the three Ser, as several chemoproteomic studies have identified YTHDF1 O-GlcNAcylation sites to be Ser196-198 (27-29). Thus we generated a YTHDF1-S196AS197FS198A (AFA) mutant. When we transfected the WT and AFA mutant into cells, the AFA mutant significantly diminished YTHDF1 O-GlcNAcylation levels (Fig. 1F), suggesting that they are the main glycosylation sites.

### Crm1 mediates the nuclearcytoplasmic shuttling of YTHDF1

To investigate the potential YTHDF1 O-GlcNAcylation functions, we first employed an immunoprecipitation-mass spectrometry (MS) analysis. Flag-YTHDF1 plasmids were transfected into cells and the lysates were immunoprecipitated with anti-Flag antibodies. Interestingly, the MS results revealed many importins and exportins (data not shown). When we did literature research, YTHDF1 was among the binding partners of exportin 1 (Crm1) in a recent proteomic study (32). Therefore, we suspect that YTHDF1 might be a nuclearcytoplasmic protein and Crm1 might mediat the process.

To test this possibility, we first assessed the association between YTHDF1 and Crm1. We also utilized KPT-330, a Crm1 inhibitor. Overexpressed YTHDF1 coIPs with Crm1, and KPT-3301 markedly reduced the interaction (Fig. 2A). Moreover, endogenous YTHDF1 interacts with Crm1 (Fig. 2B), suggesting that YTHDF1 could be a nuclearcytoplasmic protein. We then utilized the nuclear cytoplasmic fractionation assay, and fractionation results revealed that there is indeed a nuclear portion of YTHDF1 (Fig. 2C). We further adopted KPT-330 in the fractionation assay and found that KPT-330 significantly enhanced the nuclear fraction of YTHDF1 (Fig. 2D). Furthermore, in immunofluorescence staining samples, both endogeous YTHDF1 and overproduced YTHDF1 manifested significant upregulation of nuclear staining signals (Fig. 2E-F), suggesting that Crm1 could export YTHDF1 to the cytosol.

**Fig 2.**
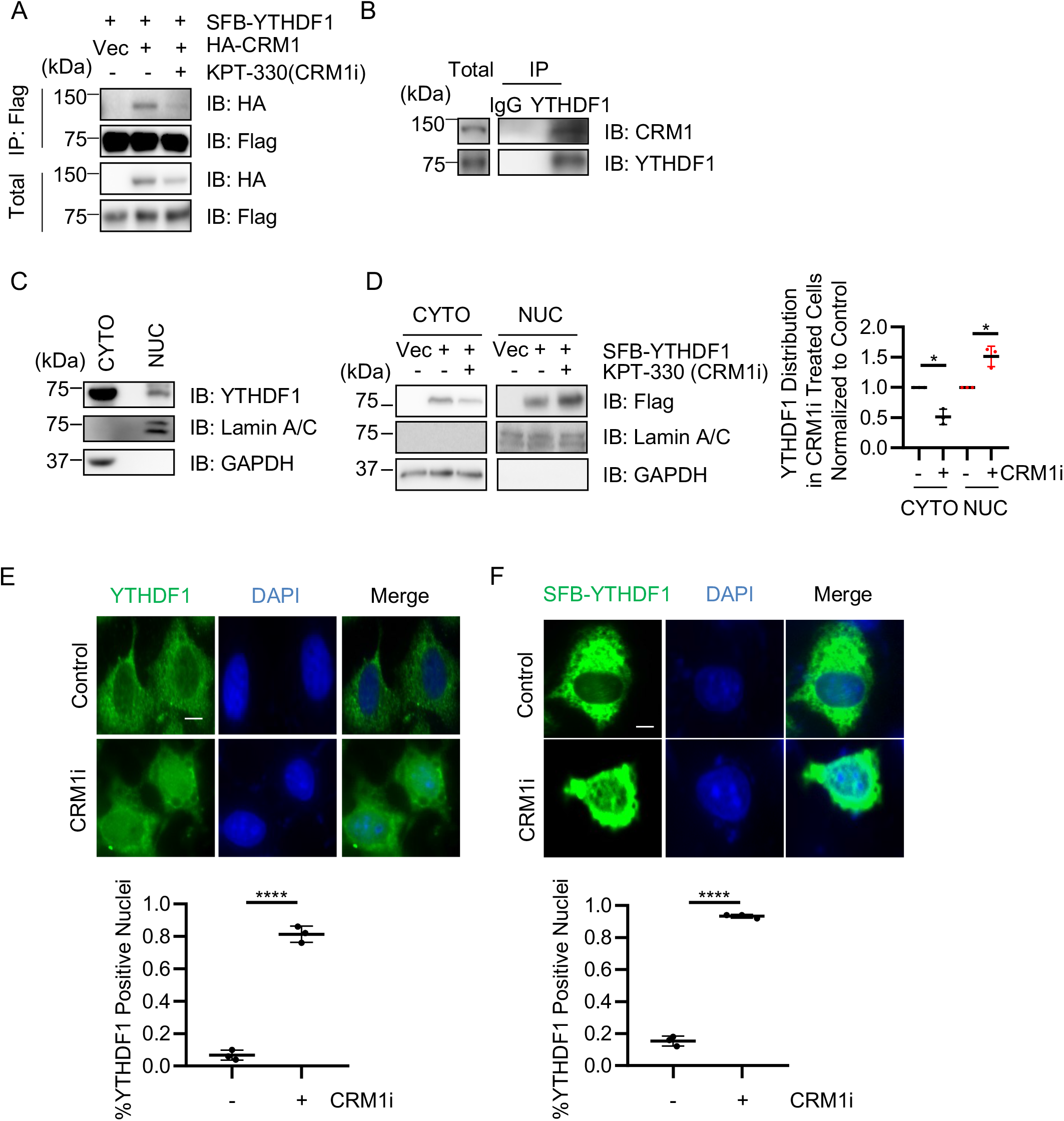
Nuclear cytoplasmic shuttling of YTHDF1 is mediated by exportin 1 (Crm1). (A) Overproduced YTHDF1 interacts with Crm1. Cells were transfected with SFB-YTHDF1 and HA-CRM1, treated or untreated with KPT-330 (Crm1 inhibitor). (B) Endogenous YTHDF1 interacts with Crm1. Cell lysates were immunoprecipitated with anti-YTHDF1 antibodies. (C) Cell lysates were subject to nuclear cytoplasmic fractionation to indicate cytosolic (CYTO) and nuclear (NUC) portions. (D) KPT-330 treatment increases nuclear YTHDF1. Cells were transfected with SFB-YTHDF1 and treated with or without KPT-330. (E-F) Indirect immunofluorescence demonstrated that KPT-330 treatment increases the nuclear localization of endogenous YTHDF1 (E) and overexpressed YTHDF1 (F). Scale bar, 10 μM. * indicates p<0.05; **** indicates p<0.0001.

### YTHDF1 O-GlcNAcylation promotes interaction with Crm1

To determine if O-GlcNAcylation plays a role in Crm1-mediated YTHDF1 nuclear shuttling, we enriched for protein O-GlcNAcylation by TMG + Glu as previously described (30). We found that O-GlcNAc enrichment increased the binding between YTHDF1 and Crm1 (Fig. 3A). We also repressed O-GlcNAcylation by an OGT inhibitor, Acetyl-5S-GlcNAc (5S-G) (33). 5S-G treatment significantly reduced the affinity between YTHDF1 and Crm1 (Fig. 3B). When 5S-G was included in the fractionation assay, the nuclear YTHDF1 is upregulated notably (Fig. 5C), suggesting that O-GlcNAcylation increases the binding between YTHDF1 and Crm1.

**Fig. 3.**
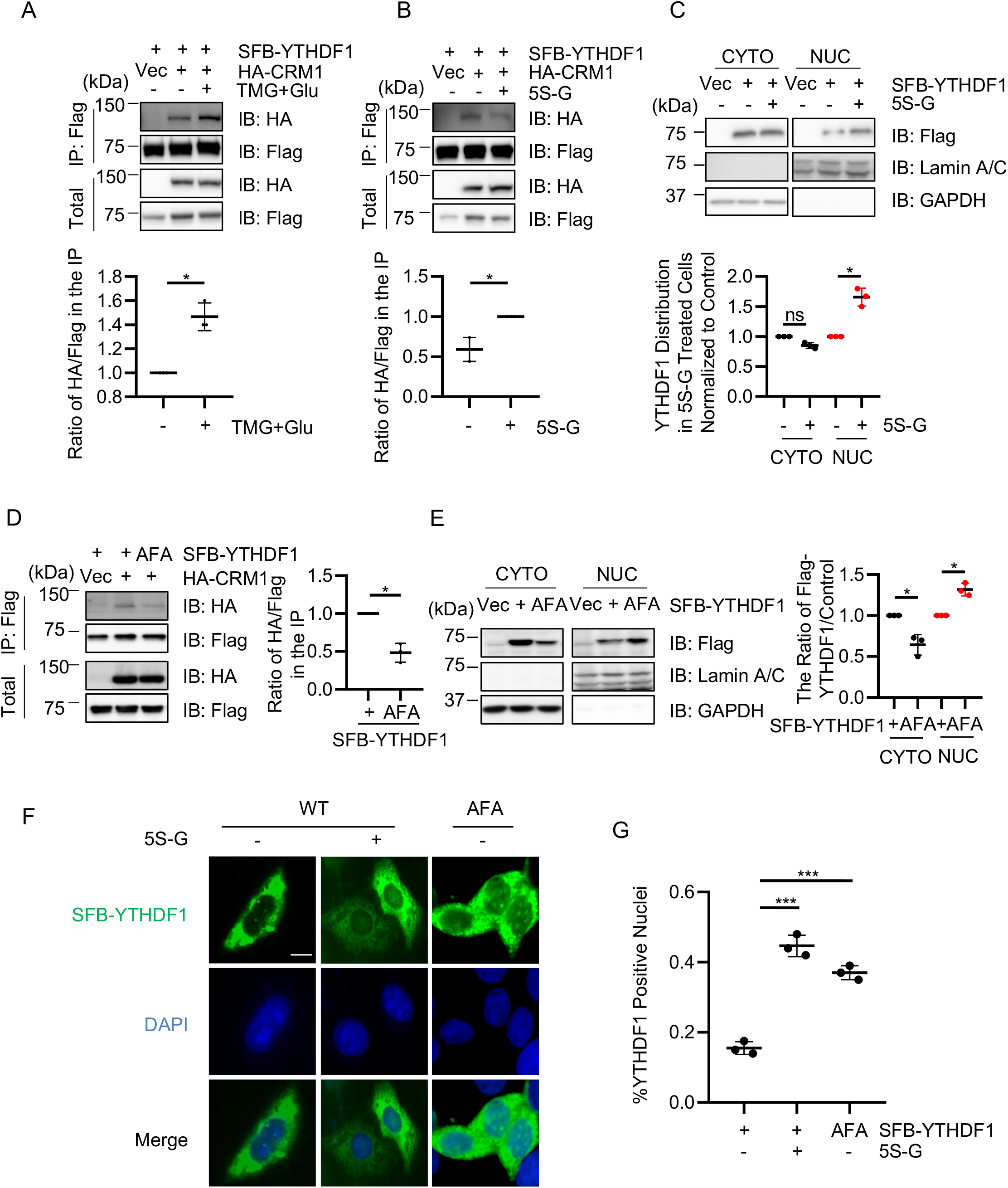
O-GlcNAcylation promotes the interaction between YTHDF1 and Crm1. (A) Cells were transfected with SFB-YTHDF1 and HA-CRM1 and enriched for O-GlcNAcylation by TMG plus glucose treatment (TMG + Glu) as previously described (30). And O-GlcNAcylation enhances the binding between YTHDF1 and Crm1. (B) Cells were transfected with SFB-YTHDF1 and HA-CRM1 and treated with the OGT inhibitor Acetyl-5S-GlcNAc (5S-G). And OGT inhibition downregulated the affinity between YTHDF1 and Crm1. (C) Cells were transfected with SFB-YTHDF1 and treated with 5S-G. Nuclear and cytoplasmic fractionation assays were carried out. OGT inhibition elevated nuclear YTHDF1. (D) Cells were transfected with HA-Crm1 together with SFB-YTHDF1-WT or -AFA plasmids. (E) Cells were transfected with SFB-YTHDF1-WT or -AFA mutants and subject to nuclear and cytoplasmic fractionation assays. (F-G) Cells were transfected with SFB-YTHDF1-WT (treated or untreated with 5S-G), or -AFA. The cells were then stained with anti-Flag antibodies and DAPI. Scale bar, 10 μM. * indicates p<0.05; *** indicates p<0.001.

Then we directly measured the effect using the AFA mutant. YTHDF1-AFA displayed marked reduction in association with Crm1 (Fig. 3D). And in the fractionation analysis, AFA again manifested much higher portion in the nucleus (Fig. 3E). Lastly, we employed fluorescent microscopy to visualize whether O-GlcNAcylation could affect YTHDF1 localization. As shown in Figure 3F-G, both 5S-G treatment and the AFA mutant enhanced nuclear YTHDF1 staining, probably by blocking its nuclear export via Crm1. These assays suggest that YTHDF1 O-GlcNAcylation promotes the binding between YTHDF1 and Crm1 and the resultant nuclear export.

### A potential Nuclear Export Signal (NES) lies in proximity to YTHDF1 O-GlcNAcylation sites

We are curious why O-GlcNAcylation has such a conspicuous effect on YTHDF1 localization and looked for potential nuclear export signals (NES) surrounding the S196S197S198 region. As NES consists of the Φ1-X(2-3)-Φ2-X(2-3)-Φ3-X-Φ4 motif (Φ: hydrophobic amino acid) (34), we found a potential NES justaposing the 196-198 Ser cluster (Fig. 4A). We mutated the corresponding hydrophobic amino acid to Ala and generated 4A (Fig. 4A), as previously described for the NES of the cyclic GMP-AMP (cGAMP) synthase (cGAS) (35). When we examined for YTHDF1-Crm1 association, the 4A mutant significantly downregulated the binding with Crm1 (Fig. 4B). In the fractionation studies, 4A also elevated nuclear YTHDF1 localization (Fig. 4C). In the immunofluorescent staining experiments, 4A also has a more prominent nuclear localization pattern compared to the control (Fig. 4D). Combined, these data suggest that O-GlcNAcylation might boost the association of the neighbouring NES with Crm1.

**Fig. 4.**
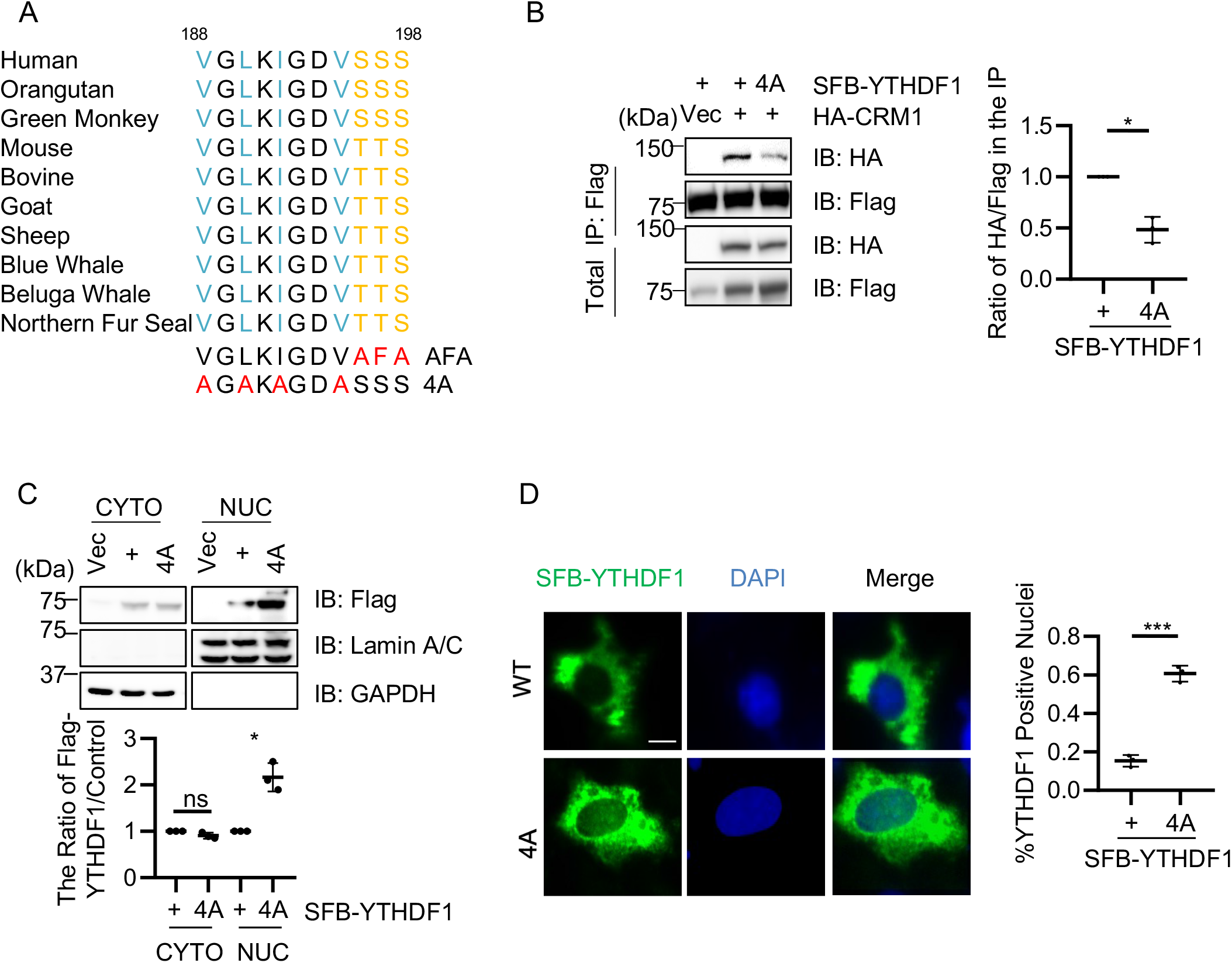
There is a potential nuclear exportin signal (NES) in proximity to O-GlcNAcylation sites. (A) Sequence alignment of the YTHDF1-WT, O-GlcNAc-deficient AFA, and the NES-deficient 4A sequences. (B) Cells were transfected with HA-Crm1, together with YTHDF1-WT and -4A plasmids. (C) Cells were transfected with SFB-YTHDF1-WT or -4A plasmids and analyzed by nuclear cytoplasmic fractionation. (D) Cells were transfected with SFB-YTHDF1 or -4A, and stained with anti-Flag antibodies and DAPI. Scale bar, 10 μM. * indicates p<0.05; *** indicates p<0.001, ns indicates non-specific.

### Molecular dynamics (MD) simulations suggest that S197 O-GlcNAcylation increases the interaction between NES and Crm1 via hydrogen bonds

We then explored deeper as why O-GlcNAcylation increases binding with Crm1. Recently a structural study focusing on the interface between NES and CRM1 found that many NESs might form hydrogen bonds with CRM1 (36), therefore we wondered whether hydrophilic O-GlcNAc could enhance the interaction by increasing hydrogen bonding. And we utilized the molecular dynamics (MD) simulation approach and began by constructing the system. Since the AlphaFold Protein Structure Database cannot well predict the NES domain of YTHDF1 (pLDDT < 50)(37,38), the ColabFold web server was used to build the initial structure of a short fragment (residues 182 - 210) including the NES domain (Fig. 5A)(39). The initial structure was further optimized for 300 ns with molecular dynamics simulations (Fig. 5B and C). The binding domain of CRM1 (residues 362 – 645, Fig.5D) was cropped from the crystal structure of the PKI NES-CRM1-RanGTP nuclear export complex (PDB ID: 3NBY)(40). The Rosetta Docking protocol (version 3.12) was applied to build the YTHDF1 NES and CRM1 complex (41-43). The NES fragment was set as the input structure with 10 Å translation and 360º rotation. One hundred poses were created after the docking process (Fig. 5E) and only two obtained reasonable relative positions (the NES domain is close to the CRM1 binding domain) (Fig. 5F). After 500 ns of MD simulations, only one complex maintained a reasonable interaction (Fig. 5G). The last frame of this complex was chosen as the initial structure for further analysis.

**Fig. 5.**
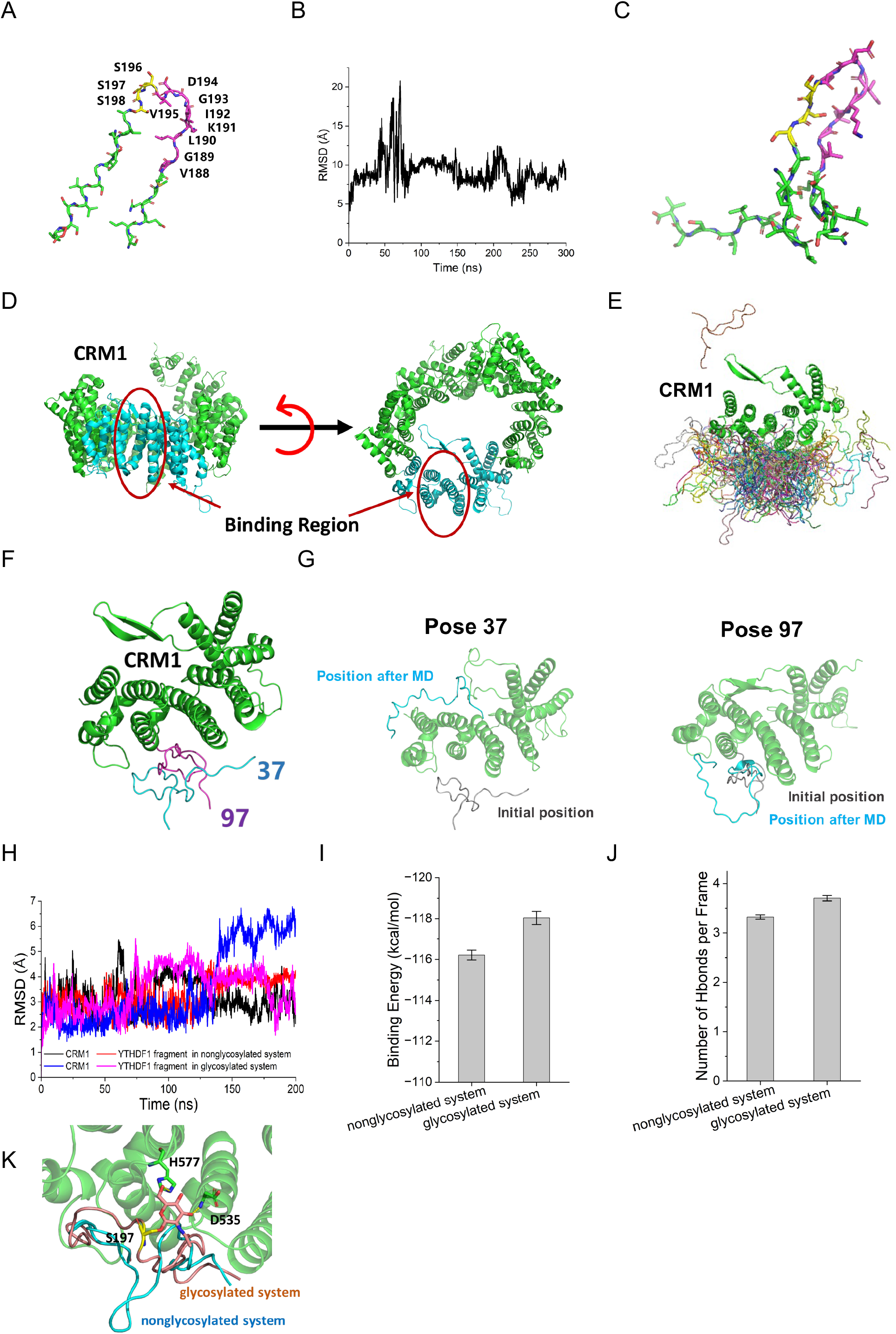
Molecular dynamics (MD) simulations suggest that S197 O-GlcNAcylation increases the interaction between NES and Crm1 via hydrogen bonds. (A) Initial structure of YTHDF1 fragment from ColabFold. The NES domain is colored in magenta and serines that could be glycosylated are colored in yellow. (B) Root-mean-square deviation (RMSD) of backbone of YTHDF1 fragments during 300 ns of MD simulation. (C) Structure of YTHDF1 fragments after optimization. (D) Structure of CRM1. The NES binding region cropped for docking is shown in cyan. (E) Docking results for CRM1 and the YTHDF1 fragment. (F). Reasonable poses (Pose 37 in cyan and Pose 97 in magenta) chosen from 100 poses. (G) Positions of YTHDF1 before and after MD simulations in Poses 37 and 97. (H) RMSDs of the backbone of CRM1 and YTHDF1 fragments in the non-glycosylated system (black for CRM1 and red for YTHDF1 fragments) and the glycosylated system (blue for CRM1 and magenta for YTHDF1 fragment) during 200 ns of MD simulations. (I) Binding energies between CRM1 and YTHDF1 fragments in the non-glycosylated and glycosylated systems. (J) Number of hydrogen bonds per frame in the non-glycosylated and glycosylated systems. (K) Detailed interaction between glycan and key residues in CRM1. The hydrogen bonds are shown in yellow dashed lines.

**Fig. 6.**
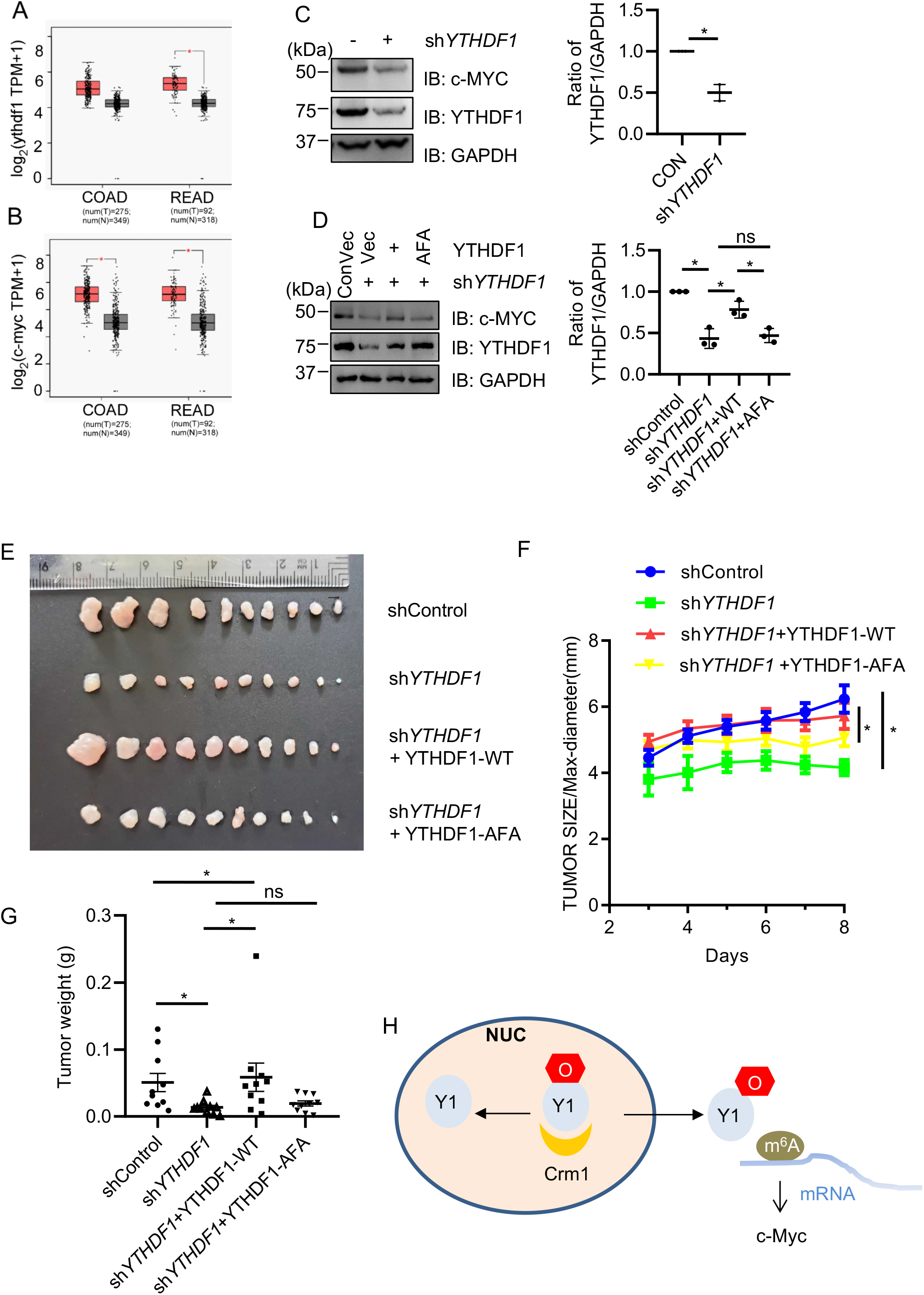
YTHDF1 O-GlcNAcylation promotes c-Myc expression in colorectal carcinoma. (A-B) YTHDF1 and c-Myc mRNA levels in colon adenocarcinoma (COAD) and rectum adenocarcinoma (READ) samples from The Cancer Genome Atlas (TCGA) database. (C) Stable YTHDF1-knockdown SW620 cell lines were generated and examined for c-Myc expression. (D) The cell lines in (C) were rescued with Flag-YTHDF1-WT, or -AFA plasmids. And cellular lysates were immunoblotted with antibodies indicated. (E-G) Xenografts in nude mice. The stable SW620 cells were injected into nude mice. The tumors were imaged after 8 days. E shows the tumor images, F shows the tumor size, and G shows the tumor weight. * indicates p<0.05. (H) A model illustrating the role of O-GlcNAc in YTHDF1 nuclear shuttling. O-GlcNAcylation of YTHDF1 at S196S197S198 will enhance the partnership between Crm1 and YTHDF1, thus promoting cytosol localization of YTHDF1 and translation of downstream target proteins, e.g., c-Myc. Such an OGT-YTHDF1-c-Myc pathway will enhance colorectal cancer.

The root-mean-square deviation (RMSD) values indicated that both systems can reach stable states in 200 ns (Fig. 5H). The trajectory of the last 100 ns was extracted for further analysis. The binding energy of glycosylated fragment to CRM1 binding domain was −118.04 ± 0.32 kcal/mol, which is lower than that of the unglycosylated fragment to the CRM1 binding domain (−116.21 ± 0.24 kcal/mol) (Fig. 5I). The number of hydrogen bonds between the fragment and CRM1 was increased when S197 was glycosylated (3.70 ± 0.06 in the glycosylated system vs. 3.32 ± 0.04 in the non-glycosylated system, Fig. 5J) because the glycan at S197 can frequently form hydrogen bonds with H577 and D535 in CRM1 to pull the NES domain to the CRM1 binding domain (Fig. 5K). Taken together, MD simulations suggest that O-GlcNAc might increase hydrogen bonding between YTHDF1 and Crm1.

### YTHDF1 O-GlcNAcylation promotes downstream target expression (e.g. c-Myc)

Recently, many YTHDF1-mediated m^6^A mRNA targets have been identified, such as the protein translation initiation factor EIF3 (16), the key Wnt receptor frizzled7 (FZD7) (15) and c-Myc (13). We focused on c-Myc, as m^6^A-modified c-Myc mRNA has been demonstrated to recruit YTHDF1 (13). We reasoned that YTHDF1 O-GlcNAcylation would promote c-Myc expression as there is more cytosolic YTHDF1. We first carried out a TCGA analysis, and found that in colon adenocarcinoma (COAD) and rectum adenocarcinoma (READ) samples, both YTHDF1 and c-Myc are overexpressed in the tumor samples (Fig. 5A-B), indicative of a positive correlation between YTHDF1 and c-Myc in colorectal cancer. We therefore generated stable YTHDF1-knockdown SW620 cells using sh*YTHDF1*, and indeed c-Myc protein levels are attenuated upon YTHDF1 downregulation (Fig. 5C). When the knockdown cells were rescued with YTHDF1-WT or -AFA plasmids, c-Myc expression is comparable to the control in the YTHDF1-WT rescued cells, but not in the -AFA rescued cells (Fig. 5D). The stable SW620 cells were then utilized in mouse xenograft experiments, and the tumor size and weight were monitored (Fig. 5E-G). As expected, the YTHDF1-WT rescued cells produced much larger tumors compared to the AFA mutants, suggesting that YTHDF1 O-GlcNAcylation promotes colorectal cancer, probably via c-Myc.

## Discussion

In this work, we first confirmed that YTHDF1 O-GlcNAcylation occurs on Ser196/197/198, then we identified that glycosylation promotes shuttling of YTHDF1 to the cytoplasm by CRM1. Consequently, cytosolic YTHDF1 will upregulate its downstream target expression (e.g., c-Myc), and then tumorigenesis will ensue.

Our work is in line with the observation that O-GlcNAcylation elevation correlates with different cancer types, such as breast cancer, prostate cancer, bladder cancer (44). In colon cancer, both O-GlcNAc and OGT abundance increased in clinical patient samples (45). Here we found that YTHDF1 O-GlcNAcylation boost the expression of c-Myc, at least in SW620 cells. In xenograft models, the O-GlcNAc-deficient YTHDF1-AFA mutants attenuated tumor progression, suggesting that OGT could regulate many more downstream substrates to promote cancer growth.

Of the many m^6^A readers, YTHDF1-3 have been considered as cytosolic proteins (46). We show here that YTHDF1 is partly localized to the nucleus, and we found a potential NES in YTHDF1. Incidentally, the NES neighbors the O-GlcNAcylation sites, suggesting that O-GlcNAcylation might promote the interaction between NES and Crm1. MD simulations suggest that the hydrophilic O-GlcNAcylation might increase the binding between NES and Crm1 through hydrogen bonding.

A great many investigations have shown that O-GlcNAcylation alters protein localization, such as pyruvate kinase M2 (PKM2) (47) and serine/arginine-rich protein kinase 2 (SRPK2) (48). PKM2 O-GlcNAcylation at Thr405/Ser406 promotes ERK-dependent phosphorylation of PKM2 at Ser37, which is required for PKM2 nuclear translocation (47,49). And PKM2-T405A/S406A attenuates interaction with importin α5(47). SRSF Protein Kinase 2 (SRPK2) is O-GlcNAcylated at Ser490/Thr492/Thr498, which is close to a nuclear localization signal (NLS) (48). And this NLS mediates SRPK2 nuclear localization by importin α (48). Indeed, a general mechanism has been proposed that at least some O-GlcNAcylated proteins are imported to the nucleus by importin α (48). Our work here suggest that maybe in some other cases, O-GlcNAcylation might shuttle the O-GlcNAcylated proteins to the cytoplasm by exportin.

The intertwined relationship between RNA and glycosylation is just emerging. Recently, a “glycoRNA”concept has been coined as small noncoding RNAs are found to be decorated with sialylated glycans (50). As far as m^6^A is concerned, many readers have been identified in O-GlcNAc chemoproteomic profiling works (25,26,29,51,52), including YTHDF1, YTHDF3 and YTHDC1. In a recent investigation from our group (https://doi.org/10.1101/2022.09.03.506498), we found that YTHDC1 O-GlcNAcylation is induced upon DNA damage and takes part in homologous recombination by enhancing binding with m^6^A mRNA. Here we show that YTHDF1 O-GlcNAcylation mediates its localization by promoting binding with exportin. We think that O-GlcNAcylation is bound to be found in many more aspects of RNA metabolism, as sweetness lies in the heart of our fellow glycobiologists.

## Materials and methods

### Cell culture, antibodies and plasmids

Cells were purchased from ATCC. OGT plasmids and antibodies were described before(53). Antibodies: YTHDF1 (Proteintech, #17479-1-AP), c-Myc (Abcam, Ab32072), Lamin A/C (CST, 2032S). YTHDF1 shRNA sequences (TRCN0000286871): 5’-CCGGCCCGAAAGAGTTTGAGTGGAACTCGAGTTCCACTCAAACTCTTTCGGGTTTTTG-3’

### Immunoprecipitation (IP) and Immunoblotting (IB) assays

IP and IB experiments were performed as described before (54). Nuclear and cytoplasmic fractionation assays were carried out as before (55). The following primary antibodies were used for IB: anti-β-actin (1:10000), anti-HA (1:1000), and anti-FLAG M2 (Sigma) (1:1000), anti-Myc (1:1000), anti-YTHDF1 (1:1000), Lamin A/C (1:1000). Peroxidase-conjugated secondary antibodies were from JacksonImmuno Research. Blotted proteins were visualized using the ECL detection system (Amersham). Signals were detected by a LAS-4000, and quantitatively analyzed by densitometry using the Multi Gauge software (Fujifilm). All western blots were repeated for at least three times.

### Cell Culture Treatment

Chemical utilization: Thiamet-G (TMG) (OGA inhibitor) at 5 μM for 24 hrs; acetyl-5S-GlcNAc (5S-G) (OGT inhibitor) was used at 100 μM (prepared at 50 mM in DMSO) for 24 hrs; KPT-300 (Crm1 inhibitor) at 5 μM for 24 hrs.

### Indirect Immunofluorescence

Indirect immunofluorescence staining was performed as described before (54). Dilutions of primary antibodies were 1:500 for mouse anti-YTHDF1, and 1:1000 for anti-Flag antibodies. Cell nuclei were stained with DAPI.

### Molecular Dynamics (MD) Simulations

The O-glycan (β-*N*-Acetyl-_D_-Glucosamine) at S197 of the YTHDF1 fragment was built using the Glycan Reader & Modeler module (56). The role of O-glycosylation in the YTHDF1 fragment interacting with CRM1 was investigated via molecular dynamics simulations with the GROMACS (version 2021.2) software package(57,58). Two systems (unglycosylated fragment and O-GlcNAcylated fragment at S197 in complex with CRM1 binding domain, respectively) were neutralized and solvated by 150 mM KCl and TIP3P water molecules. The systems were minimized and equilibrated using the default equilibration inputs from the CHARMM-GUI webserver(59) with the CHARMM36m force field (60,61). In brief, the systems were equilibrated in the isothermal-isobaric (NPT) ensemble for 200 ns. The pressure was set at 1 atm maintained by the Parrinello-Rahman barostat (62) and the temperature was maintained at 310.15 K with the Nosé–Hoover thermostat(63). Periodic boundary conditions were applied throughout the simulations. The SHAKE algorithm was used to constrain all bonds with hydrogen atoms(64). The particle-mesh Ewald (PME) summation method was applied to treat long-range electrostatic interactions (65).

Analysis of MD trajectory data was performed through MDAnalysis (66). The binding energy (enthalpy) and per-residue energy contributions were calculated by the molecular mechanics/Poisson-Boltzmann (generalized-Born) surface area method with the gmx_MMPBSA tool (67,68). The interactions between the YTHDF1 fragment and the CRM1 binding domain were displayed by PyMol (69).

### Mouse Xenograft

1 × 10^6^ control, YTHDF1 shRNA, YTHDF1 shRNA; YTHDF1-WT or YTHDF1 shRNA;YTHDF1-AFA stable SW620 cells were resuspended in Matrigel (Corning) and then injected into the flanks of nude mice (4-6 weeks old). The tumor volumes were measured from day 3 to 9 after injection. At 9 days after the injection, tumors were dissected. The mice were obtained from the Animal Research and Resource Center, Yunnan University.

{Certification NO. SCXK(Dian)K2021-0001}. All animal work procedures were approved by the Animal Care Committee of the Yunnan University (Kunming, China).

## Data Availability Statement

All data are contained within the manuscript.

## Abbreviation

(TMG): Thiamet-G
(MS): Mass spectrometry
(IP): Immunoprecipitation
(IB): Immunoblotting
(5S-G): Acetyl-5S-GlcNAc
(O-GlcNAc): O-linked β-N-acetylglucosamine
(OGT): O-GlcNAc transferase
(m^6^A): mRNA N^6^-methyladenosine
(ETD): electron transfer dissociation

## Acknowledgements

We thank Dr. Xing Chen (Peking Univ.) for reagents, and Dr. Qing Chang for providing facility support at the Protein Preparation and Characterization Platform of Tsinghua University Technology Center for Protein Research. This work was supported by the National Natural Science Foundation of China (NSFC) fund (31872720 and 32271285), R & D Program of Beijing Municipal Education Commission (KZ202210028043) to Jing L. and NSFC (82273460) fund and the Yunnan Applied Basic Research Projects (202101AV070002 and 2019FY003030) to J. S.

